# Phasic and tonic serotonin modulate alarm reactions and post-exposure behavior in zebrafish

**DOI:** 10.1101/827824

**Authors:** Monica Lima-Maximino, Maryana Pereira Pyterson, Rhayra Xavier do Carmo Silva, Gabriela Cristini Vidal Gomes, Sueslene Prado Rocha, Anderson Manoel Herculano, Denis Broock Rosemberg, Caio Maximino

## Abstract

Current theories on the role of serotonin (5-HT) in vertebrate defensive behavior suggest that this monoamine increases anxiety but decreases fear, by acting at different levels of the neuroaxis. This paradoxical, dual role of 5-HT suggests that a serotonergic tone inhibits fear responses, while an acute increase in 5-HT would produce anxiety-like behavior. However, so far no evidence for a serotonergic tone has been found. Using zebrafish alarm responses, we investigate the participation of phasic and tonic 5-HT levels in fear-like behavior, as well as in behavior after stimulation. Conspecific alarm substance (CAS) increased bottom-dwelling and erratic swimming, and animals transferred to a novel environment after CAS exposure (post-exposure behavior) showed increased bottom-dwelling and freezing. Clonazepam blocked CAS effects during and after exposure. Acute fluoxetine dose-dependently decreased fear-like behavior, but increased post-exposure freezing. Metergoline had no effect on fear-like behavior, but blocked the effects of CAS on post-exposure behavior; similar effects were observed with pCPA. Finally, CAS was shown to decrease the activity of monoamine oxidase in the zebrafish brain after exposure. These results suggest that phasic and tonic serotonin encode an aversive expectation value, switching behavior towards cautious exploration/risk assessment/anxiety when the aversive stimulus is no longer present.

## 1. Introduction

The neurocircuitry of defensive reactions involves regulation by a plethora of neuromodulators, including monoamines and peptides (Maximino 2012). In vertebrates, the monoamine serotonin (5-HT) is produced in specific brain nuclei, including the raphe, and is thought to inhibit fear/escape responses to proximate threat by acting on more caudal structures of the aversive brain system (Paul *et al.* 2014; Deakin and Graeff 1991; Maximino 2012). This response appears to be dependent on the specific brain region in which serotonin acts, as well as on the receptor that is activated. For example, in the rodent periaqueductal gray, the activation of 5-HT_1A_ and 5-HT_2_-type receptors inhibit fear responses, while in amygdaloid nuclei the activation of 5-HT_2_- and 5-HT_3_-type receptors increase anxiety-like responses (Guimarães *et al.* 2008; Paul *et al.* 2014; Hale and Lowry 2011). There is also evidence for a serotonergic “tone” inhibiting anxiety, since antagonists usually inhibit anxiety-like responses in animal models; however, antagonists do not appear to modify fear-like responses, suggesting that phasic, not tonic, serotonin is involved in fear. For example, 5-HT levels do not change in the basolateral amygdala or in the dorsal periaqueductal gray during chemical stimulation of this latter structure in rats (Zanoveli *et al.* 2009), a manipulation that induces panic-like responses (Brandão *et al.* 2008). This suggests that the inhibitory role of serotonin in fear functions as a “switching” signal: as the threatening stimulus ceases, serotonin is released, inhibiting the fear reactions that are now non-adaptive, and initiating careful exploration and risk assessment responses to ensure that the threat is actually over.

In non-mammalian vertebrates, including teleost fish, 5-HT is produced in additional brain regions, including pretectal and hypothalamic populations (Herculano and Maximino 2014). There is some evidence that 5-HTergic neurons innervate areas of the teleostean brain which participate in defensive behavior, including prosencephalic and mesencephalic regions (do Carmo Silva *et al.* 2018a). A role for 5-HT in modulating fish defensive behavior has been demonstrated before: in zebrafish, 5-HT_1A_ and 5-HT_1B_ receptor antagonists decrease anxiety-like behavior (Maximino *et al.* 2013; Nowicki *et al.* 2014; Herculano *et al.* 2015; Maximino *et al.* 2015), while 5-HT_2_- and 5-HT_3_-type antagonists increase it (Nowicki *et al.* 2014). Little is known, however, of the modulation of fear-like responses. Microinjection of the serotonergic neurotoxin 5,7-dihydroxytryptamine (5,7-DHT) in the telencephalon of zebrafish, destroying most serotonergic innervation in regions associated with aversive learning, impairs the acquisition of active avoidance (Amo *et al.* 2014), suggesting that serotonin encodes an aversive expectation value.

In zebrafish and other Actinopterygian fish, specialized club cells in the skin produce a substance (conspecific alarm substance, CAS) that, when the skin is damaged, is dispersed in the water, signaling to conspecifics a potential threat (von Frisch 1941; Hüttel 1941; von Frisch 1938; Døving and Lastein 2009; Maximino *et al.* 2019). CAS induces defensive behavior in zebrafish, including increased bottom-dwelling, erratic swimming, and freezing (Maximino *et al.* 2019; Egan *et al.* 2009; Speedie and Gerlai 2008; Maximino *et al.* 2014). These responses have been exploited as a model system to study fear in more basal vertebrates (Maximino *et al.* 2019; Jesuthasan and Mathuru 2008).

The serotonergic system has also been implicated in some of these behavioral functions. CAS increases extracellular serotonin levels (Maximino *et al.* 2014) and inhibits monoamine oxidase activity (Quadros *et al.* 2018) in the zebrafish brain after exposure. Zebrafish exposed to CAS show increased anxiety-like behavior in the light/dark test after exposure (i.e., when the substance is no longer present), an effect that is blocked by fluoxetine but not by the 5-HT_1A_ receptor antagonist WAY 100,635 (Maximino *et al.* 2014). Interestingly, WAY 100,635 blocked the analgesic effects of CAS in zebrafish (Maximino *et al.* 2014), suggesting that this receptor participates in some, but not all, neurobehavioral responses to threatening stimuli. While WAY 100,635 was not able to alter anxiety-like behavior *after* exposure, the drug blocked the increased geotaxis *during* CAS exposure, both in the first minutes of exposure and in the last minutes (Nathan *et al.* 2015). Blocking 5-HT_2_-type receptors with methysergide did not affect these responses, except at a sedative dose (Nathan *et al.* 2015). These results are difficult to interpret, but suggest that CAS increases serotonergic activity after exposure, and that a serotonergic tone on the 5-HT_1A_ receptor is involved in behavioral switching after exposure – that is, when the threat is no longer present, and risk assessment begins; whether this is true for behavioral responses *during* exposure – that is, when the threat is present – is unknown.

The present paper investigated whether phasic and tonic serotonin participates in the alarm response in zebrafish during and after exposure. Our results reinforce the idea that behavior during CAS exposure is qualitatively different from behavior after CAS exposure. We also show that behavior in both contexts are differentially sensitive to clonazepam, a high potency benzodizepine commonly used in the clinical management of panic disorder (Caldirola *et al.* 2016). Increasing serotonin levels by treating zebrafish with acute fluoxetine blocked the effects of CAS during and after exposure, but blocking serotonin receptors with metergoline, or blocking serotonin synthesis with *p*CPA, produced an effect only after exposure. Finally, we show that CAS inhibited monoamine oxidase activity in the brain. Results are discussed in terms of the putative role of serotonin in an homeostatic “neurobehavioral switch” in the absence of threat after predatory risk.

## 2. Methods

### 2.1. Animals, housing, and baseline conditions

435 zebrafish (*Danio rerio*) from the longfin phenotype were used in the present experiments; details for sample size calculations can be found on each experimental section, below (Figure 1). Outbred populations were used due to their increased genetic variability, decreasing the effects of random genetic drift that could lead to the development of uniquely heritable traits (Parra *et al.* 2009; Speedie and Gerlai 2008). The populations used in the present experiments are expected to better represent the natural populations in the wild. Animals were bought from a commercial vendor (Fernando Peixes, Belém/PA) and arrived in the laboratory with an approximate age of 3 months (standard length = 13.2 ± 1.4 mm), and were quarantined for two weeks; the experiment began when animals had an approximate age of 4 months (standard length = 23.0 ± 3.2 mm). Animals were kept in mixed-sex tanks during acclimation, with an approximate ratio of 50-50 males to females (confirmed by body morphology). The breeder was licensed for aquaculture under Ibama’s (Instituto Brasileiro do Meio Ambiente e dos Recursos Naturais Renováveis) Resolution 95/1993. Animals were group-housed in 40 L tanks, with a maximum density of 25 fish per tank, for at least 2 weeks before experiments begun. Tanks were filled with non-chlorinated water at room temperature (28 °C) and a pH of 7.0-8.0. Lighting was provided by fluorescent lamps in a cycle of 14-10 hours (LD), according to standards of care for zebrafish (Lawrence, 2007). Water quality parameters were as follows: pH 7.0-8.0; hardness 100-150 mg/L CaCO3; dissolved oxygen 7.5-8.0 mg/L; ammonia and nitrite < 0.001 ppm. Potential suffering of animals was minimized by controlling for the aforementioned environmental variables and scoring humane endpoints (clinical signs, behavioral changes, bacteriological status), following Brazilian legislation (Conselho Nacional de Controle de Experimentação Animal - CONCEA 2017). Animals were used for only one experiment and in a single behavioral test, to reduce interference from apparatus exposure. Experiments were approved by UEPA’s IACUC under protocol 06/18.

**Figure 1.**
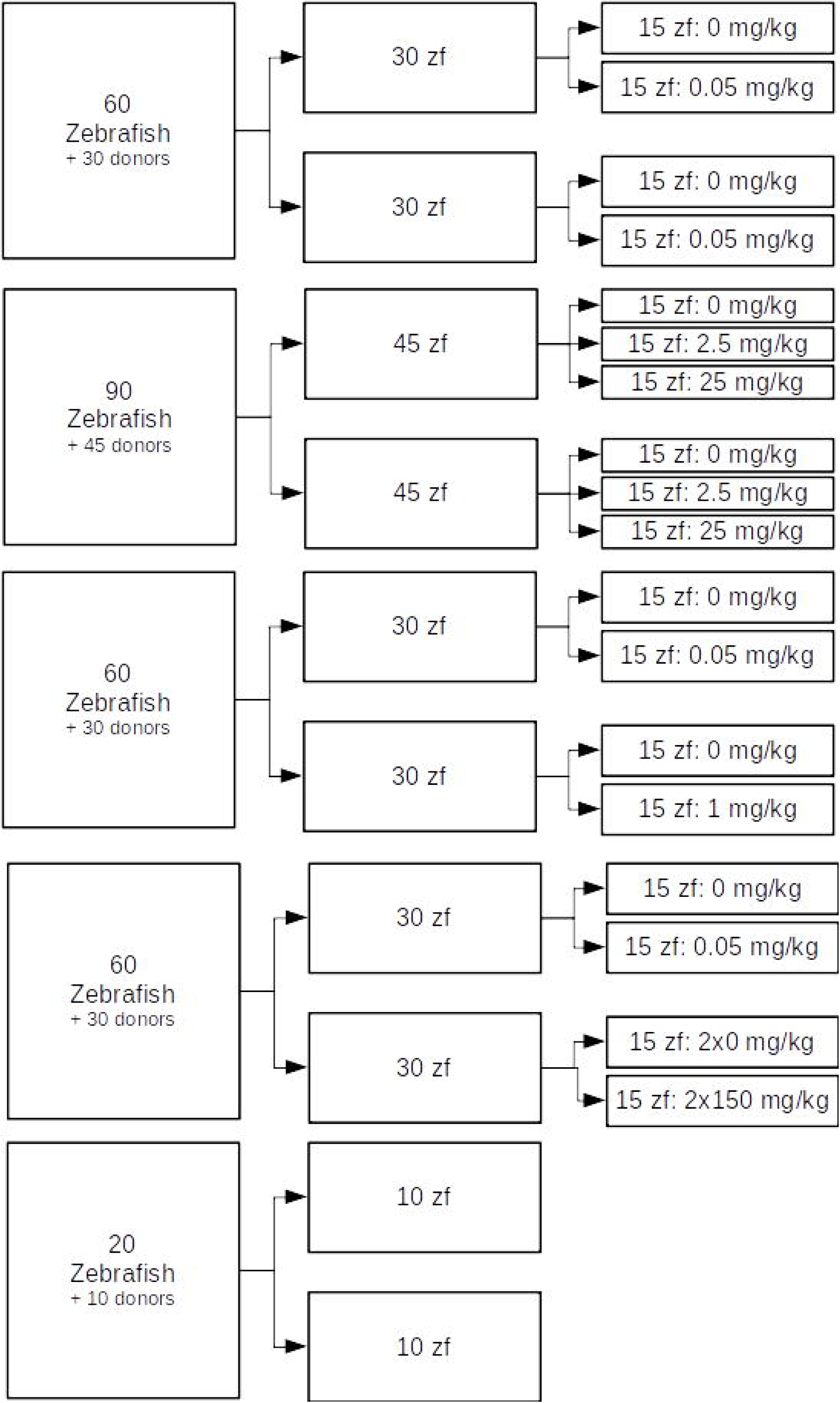
Experimental design and sample sizes for each experiment. In the right-most boxes, “donors” refer to animals which were sacrificed and used to produce alarm substance, and therefore not used as subjects.

### 2.2. Alarm substance extraction

CAS was extracted at a ratio of 1 donor fish for 10 ml distilled water. A detailed protocol for extraction can be found at protocols.io (Silva, Rocha, Lima-Maximino, & Maximino, 2018; https://dx.doi.org/10.17504/protocols.io.tr3em8n). Briefly, a donor fish was cold-anesthetized and euthanized, and 15 shallow cuts were made on the side of its trunk to lesion club cells. The cuts were washed with 10 mL distilled water, and 7 mL of the eluate was reserved as 1 unit CAS.

### 2.3. General experimental design

After the onset of drug effects (see details below for each drug), animals were individually transferred to a 1.5 L tank (12 cm X 12 cm x C cm, w X l X h), filled with system water, and left to acclimate for 3 min. Filming was started, and animals were exposed to either 7 mL distilled water (CTRL groups) or 7 mL (1 unit) alarm substance (CAS groups). Exposure was made by slowly pouring the substance on the water from the top. Animals were then left undisturbed as filming continued for 6 min; this was termed “alarm reaction”. The animal was then transferred to a tank with 500 mL mineral water for a 1 min “washout” period, to remove potential residues from the alarm substance. After this period, the animal was transferred to a 5 L tank (A cm X 24 cm X 22 cm, w X l X h) and freely explored for 6 min, during which its behavior was recorded; this was termed “post-exposure behavior” (Figure 2A). Tanks for both stages were differently shaped to increase the novelty of the second environment, a variable that is important to induce an anxiety-like “diving” response in animals not exposed to CAS (Bencan *et al.* 2009). Light levels above the tanks were measured using a handheld light meter, and ranged from 251 to 280 lumens (coefficient of variation = 3.399% between subjects) In all experiments, the following variables were recorded:

- Time spent on the bottom third of the tank (s) [Primary outcome]
- Time spent on the top third of the tank (s) [Secondary outcome]
- Absolute turn angle (equivalent to erratic swimming) [Secondary outcome]
- Freezing: duration of complete movement cessation, defined as speed lower than 0.5 cm/s. [Secondary outcome]
- Swimming speed (cm/s) [Secondary outcome]

**Figure 2.**
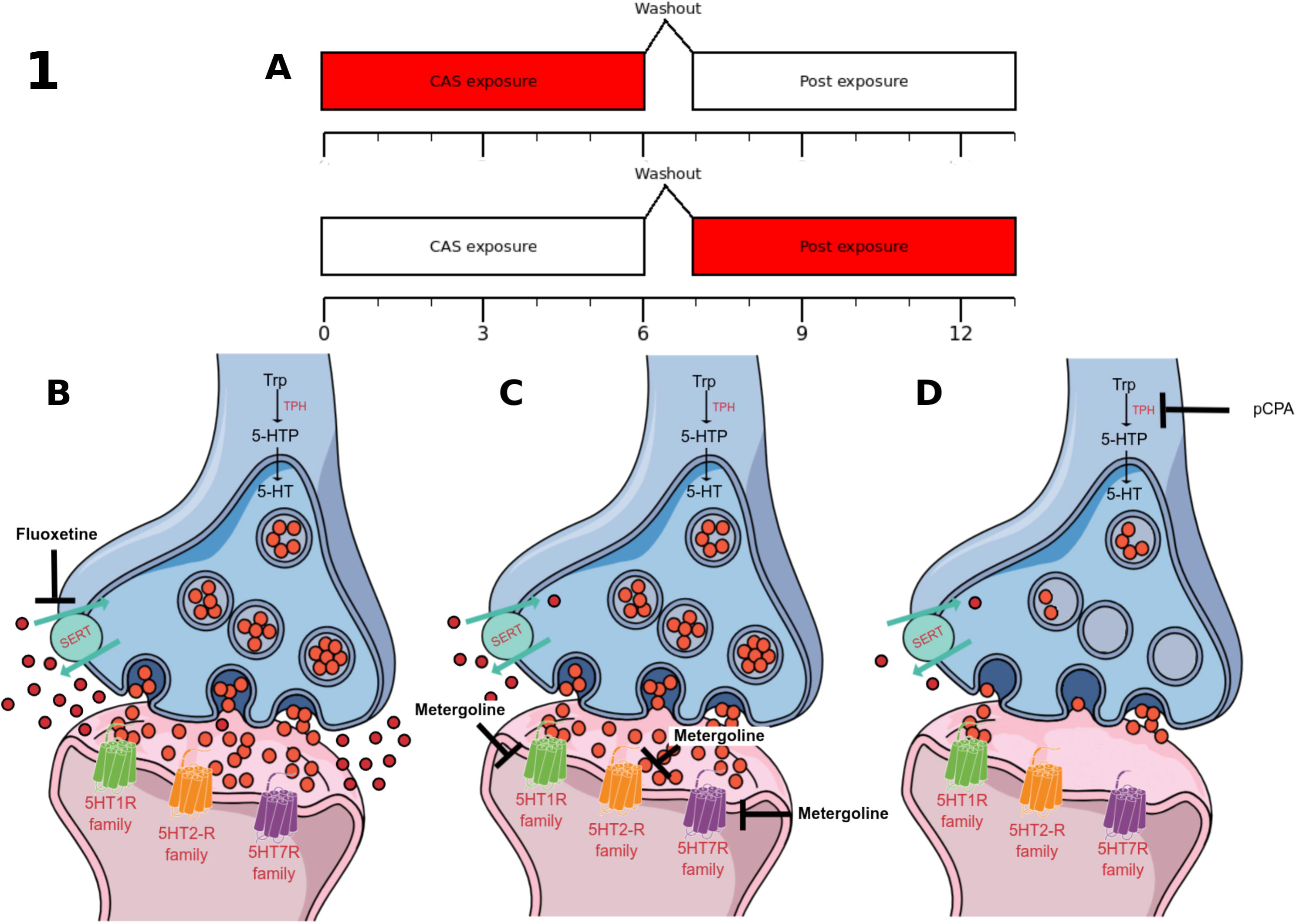
(A) Time-course of the experiments, with behavioral observations during two 6-min blocks, “CAS exposure” and “Post exposure”, separated with a 1-min washout period. The boxes in red indicate the moment that the observation is made in each block. (B-D) Representation of the synaptic effects of the pharmacological manipulations: acute fluoxetine (B) is expected to increase synaptic and extra-synaptic serotonin levels, while metergoline (C) is expected to block receptors from the 5-HT_1_, 5-HT_2_, and 5-HT_7_ family; pCPA (D) is expected to decrease synaptic and extra-synaptic serotonin levels.

Variables were extracted by automated video tracking, using the software TheRealFishTracker v. 0.4.0 (http://www.dgp.toronto.edu/~mccrae/projects/FishTracker/), running on a Windows platform. Animals were randomly allocated to groups using a random number generator (http://www.jerrydallal.com/random/random_block_size_r.htm), with each subject randomized to a single treatment using random permuted blocks. One PI attributed a random letter to treatment (e.g., “A” for CTRL, “B” for CAS) and a random integer for drug dose (e.g., “1” for 1 mg/kg, “2” for 0 mg/kg [vehicle]), and combinations for letters and integers were randomized. For each experiment, animals were treated and tested in the order of allocation (i.e., randomly). In all experiments, experimenters and data analysts were blinded to drugs and treatment by using coded vials (with the same code used for randomization); blinding was removed only after data analysis. Experiments were always run between 08:00AM and 02:00 PM. After experiments, animals were sacrificed by prolonged bath in ice-cold water (< 12 °C), followed by spinal transection (Matthews and Varga 2011).

### 2.4. Quality control

#### Exclusion criteria

With the exception of outlier exclusion (described in 2.5.3), no exclusion criteria were predetermined.

#### Behavioral data

Quality control of samples was maintained by periodic assessment of water quality and health parameters. All experimenters were trained in the behavioral methods before experiments; training included observation of all experiments by a PI (CM or MGL) on at least two occasions. After these observations, each trainee performed two mock experiments, on a single subject each, while being observed by the PI. All protocols were reviewed by all PIs, and are publicly available. Behavioral records were reviewed by at least one PI for administration/scoring accuracy, in order to ensure adherence to protocols and consistency across tests.

#### Biochemical data

All experimenters were trained in the analytical method before experiments. Quality control was achieved periodically using Levey-Jennings charts for known concentrations of kynuramine, adopting a 1_2S_ rule.

### 2.5. Experiments 1-4: Effects of clonazepam and serotonergic drugs on alarm reaction and post-exposure behavior

#### 2.5.1. Sample size calculations

Sample size calculations were based on a power analysis for a 2-way ANOVA with interaction effects, with α = 0.05, β = 0.8, and expected effect size *f* = 0.25 for each independent variable (IV); Effect sizes used for estimating sample sizes were based on the range of effects observed after pharmacological manipulations on zebrafish anxiety-like behavior in a metanalysis (Kysil et al., 2017). Sample size of 15 animals/group was established. Thus, a total of 270 animals were used for experiments, and another 135 animals were used to produce CAS. The distribution of samples through groups can be found in Figure 1.

#### 2.5.2. Drugs and treatments

Clonazepam (CAS #1622-61-3) was bought from Roche on 2018, and dissolved in Cortland’s salt solution (NaCl 124.1 mM, KCl 5.1 mM, Na_2_HPO_4_ 2.9 mM, MgSO_4_ 1.9 mM, CaCl_2_ 1.4 mM, NaHCO_3_ 11.9 mM, Polyvinylpyrrolidone 4%, 1,000 USP units Heparin; Wolf 1963). Clonazepam, as a high-potency benzodiazepine widely used in treating panic disorder (Cloos 2005; Caldirola *et al.* 2017), is expected to decrease fear-like responses to CAS, and therefore used as a positive control. Fluoxetine is expected to acutely increase serotonin levels in the synapse (Figure 2B). Fluoxetine hydrochloride (CAS #54910-89-3) was bought from Libbs on 2017, and dissolved in Cortland’s salt solution. Metergoline is expected to block 5-HT receptors from the 5-HT_1_, 5-HT_2_, and 5-HT_7_ families (Figure 2C). Metergoline (CAS #17692-51-2) was bought from Virbac on 2017, and dissolved in Cortland’s salt solution. pCPA is expected block 5-HT synthesis, therefore greatly reducing serotonergic tone on all receptors (Figure 2D). 4-chloro-DL-phenylalanine (pCPA; CAS #7424-00-2) was bought from Sigma-Aldrich (C6506) on 2018, and dissolved in 10% DMSO. For Experiment 1, animals were injected intraperitoneally with either vehicle (Cortland’s salt solution) or clonazepam (0.05 mg/kg; Maximino, Silva, Gouveia Jr., & Herculano, 2011). For Experiment 2, animals were injected intraperitoneally with vehicle (Cortland’s salt solution) or fluoxetine (2.5 or 25 /kg; Maximino et al. 2014). For Experiment 3, animals were injected intraperitoneally with vehicle (Cortland’s salt solution) or metergoline (1 mg/kg; Pimentel et al. 2019). For Experiment 4, animals were injected intraperitoneally with either vehicle (DMSO) or pCPA (one injection of 150 mg/kg/day for 2 days, followed by 24 h without treatment; Curzon et al. 1978). Injections were made according to the protocol proposed by Kinkel et al. (2010); briefly, animals were cold-anesthetized and transferred to a sponge-based surgical bed, in which injection was made. Injections were made using a microsyringe (Hamilton® 701N syringe, needle size 26 gauge at cone tip), with total volumes of injection ranging from 4.81 to 5.05 µL. Cold-anesthesia has been shown to produce satisfactory results in zebrafish, with faster recovery and less animal loss than commonly used anesthetics such as MS-222 (Matthews and Varga 2011). The sponge allowed gill perfusion to be kept, minimizing suffering. 20 min after recovery, animals were subjected to CAS or water.

#### 2.5.3. Statistical analysis

Outliers were removed based on median absolute differences (MADs), using time on bottom as main endpoint; values were removed when they were higher or lower than 3 MADs around the median (Leys *et al.* 2013), and the number of outliers was reported in the results. Differences between groups were analyzed using two-way analyses of variance (ANOVAs) with robust estimators on Huber’s M-estimators, using the R package ‘rcompanion’ (Mangiafico 2017; https://cran.r-project.org/package=rcompanion). Normality was not assumed, and thus no specific test for normality was performed; however, this type of analysis is resistant to deviations from the assumptions of the traditional ordinary-least-squares ANOVA, and are robust to outliers, thus being insensitive to distributional assumptions (such as normality)(Huber 1981). Behavioral variables were included as outcomes, with treatment and drug used as independent variables; interaction between IVs was assessed as the most important predictor. P-values were adjusted for the false discovery rate.

### 2.6. Experiment 5: Effects of CAS on monoamine oxidase activity

#### 2.6.1. Sample size

Based on a power analysis for two-sample unpaired t-test. α = 0.05, power = 0.8, and expected effect size *d* = 1.5, a sample size of 10 animals/group was established. Thus, a total of 20 animals were used for experiments, and 10 more used to produce CAS. The distribution of samples through groups can be found in Figure 1.

#### 2.6.2. Methods

z-MAO activity was determined as reported previously (Quadros *et al.* 2018). Two zebrafish brains were pooled per sample and homogenized in 0.5 mL of buffer solution containing 16.8 mM Na_2_HPO_4_ and 10.6 mM KH_2_PO_4_, pH 7.4, isotonized with sucrose. Samples (n = 10 per group) were centrifuged at 1.000 x g for 5 min, and the supernatants were kept on ice for the experiments. Protein samples (approximately 100 µg) were mixed with 460 µL of assay buffer (168 mM Na_2_HPO_4_ and 10.6 mM KH_2_PO_4_, pH 7.4, isotonized with KCl) and preincubated at 37°C for 5 min. The reaction started by adding 110 µM kynuramine hydrobromide in a final volume of 700 µL, and was stopped 30 min later with 300 µL 10% trichloroacetic acid. Reaction products were further centrifuged at 16.000 x g for 5 min and supernatants (800 µL) were mixed with 1M NaOH (1 mL). Fluorescence was measured using excitation at 315 nm and emission at 380 nm. Product formation (4-hydroxyquinoline) was estimated and enzyme activity was expressed as expressed as nmol 4-OH quinoline/min/mg protein.

#### 2.6.3. Statistical analysis

Data were analyzed with an asymptotic general independence test, using the R package ‘coin’ (Hothorn et al. 2006; https://cran.r-project.org/package=coin).

### 2.7. Open science practices

Experiments were formally preregistered at Open Science Framework (https://doi.org/10.17605/OSF.IO/QM3PX). Data packages and analysis scripts for all experiments can be found at a GitHub repository (https://github.com/lanec-unifesspa/5-HT-CAS). Preprints for the manuscript can be found at bioRxiv (https://doi.org/10.1101/827824).

#### 2.7.1. Changes from pre-registration

During pre-registration, we proposed to use manual recording of behavioral variables. During experiments, we decided to use automated tracking, due to the higher availability of open source software; automated tracking allows for a better reproducibility and precision in measures but, as a trade-off, some measurements were not possible. In the present experiments, the software TheRealFishTracker (v. 0.4.0, for Windows; http://www.dgp.toronto.edu/~mccrae/projects/FishTracker/) was used. Moreover, during pre-registration we proposed to also analyze melanophore responses to CAS; this data will appear in a separate paper, on behavioral and physiological aspects of the alarm reaction, and the data is available at a GitHub repository (https://github.com/lanec-unifesspa/5-HT-CAS/tree/master/data/melanophore). Finally, the following doses were changed from pre-registration: clonazepam was reduced to 0.05 mg/kg, to avoid unwanted sedation; a second dose of fluoxetine was added to approach the range in which fluoxetine blocks fear conditioning at a high shock intensity (Santos *et al.* 2006). Due to a problem with solubility, the pCPA dose was reduced to 150 mg/kg, a dose that has been shown to reduce serotonin levels in the rat brain by about 50%, and to block the release of 5-HT elicited by electrical stimulation of the raphe (Curzon *et al.* 1978).

## 3. Results

### 3.1. Experiment 1

One outlier was removed from the group in which animals were exposed to CAS and injected with vehicle; and one outlier was removed from the group in which were exposed to water and injected with vehicle in Experiment 1. During CAS exposure, significant effects of treatment (*p* = 0.0006), dose (*p* = 0.0044, and interaction (*p* = 0.0006) were found for time on top (Figure 3A); post-hoc tests suggested that CAS decreased time on top (adjusted *p* = 3.91·10^-5^), and CLZ blocked this effect. Significant effects of treatment (*p* = 0.0466), dose (*p* = 0.0002), and interaction (*p* < 0.0001) were found for time on bottom (Figure 3B); CAS increased time on bottom (adjusted *p* =9.44·10^-6^), and CLZ blocked this effect. Main effects of treatment (*p* = 0.0004) and dose (*p* = 0.00038), as well as an interaction effect (*p* = 0.00042), were found for absolute turn angle (Figure 3C); CAS increased absolute turn angle (adjusted *p* = 0.00033), and CLZ blocked this effect. Main effects of treatment (*p* = 0.00028) and dose (*p* = 0.0031), as well as an interaction effect (*p* = 0.0021), were found for freezing (Figure 3D); again, CAS increased freezing (adjusted *p* = 1.26·10^-6^), and CLZ blocked this effect. Main effects were found for treatment (*p* = 0.04) and dose (*p* = 0.02) for speed; however, post-hoc tests failed to uncover differences between groups (Figure 3E).

**Figure 3.**
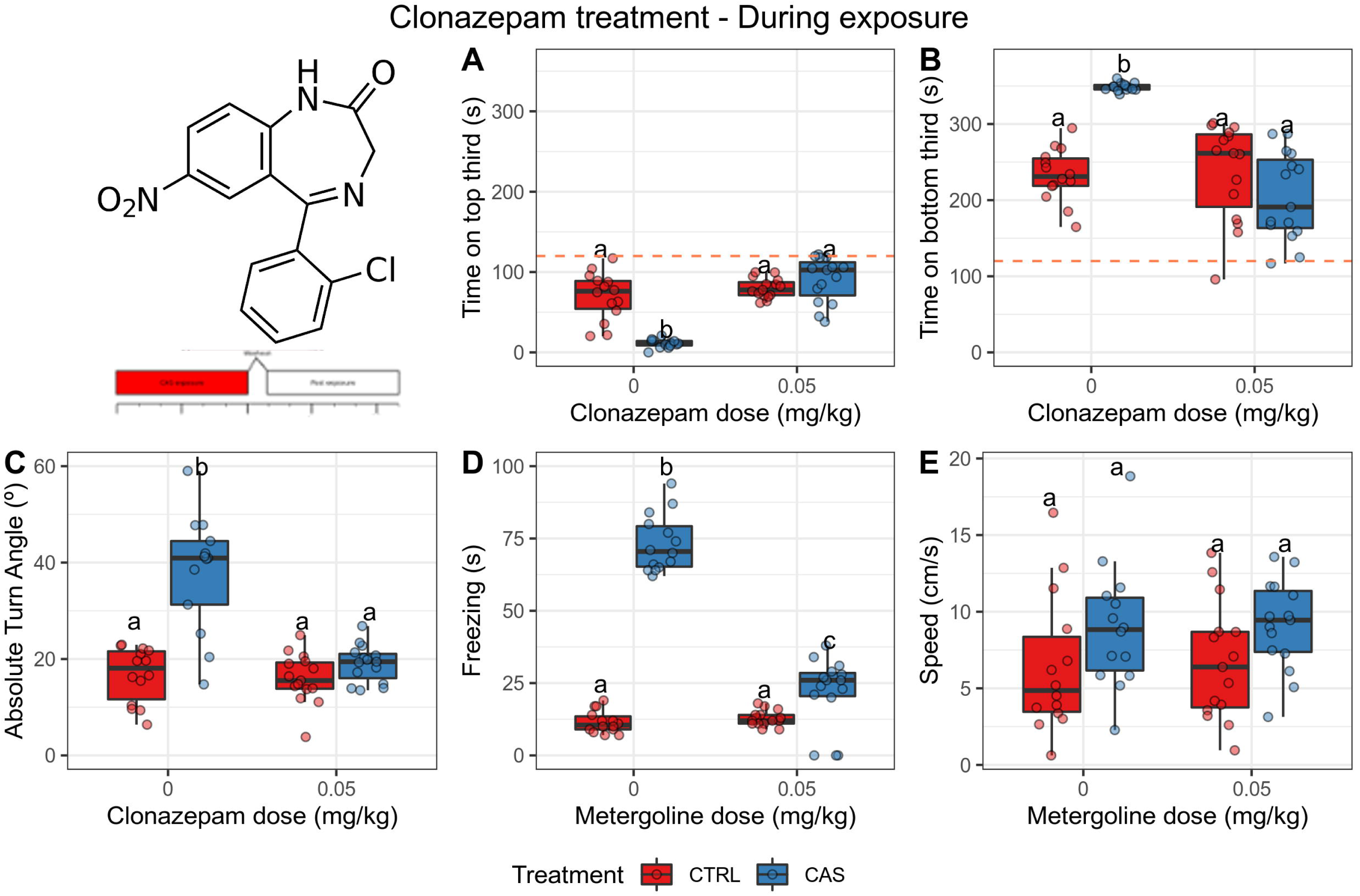
**Clonazepam (0.05 mg/kg) blocks all CAS-elicited increases in defensive behvavior during exposure.** (A) Time spent on the top third of the tank; (B) Time spent on the bottom third of the tank; (C) Absolute turn angle; (D) Freezing; (E) Swimming speed. Different letters represent statistical differences at the *p* < 0.05 level; similar letters indicate lack of statistically significant differences. Data are presented as individual data points (dots) superimposed over the median ± interquartile ranges. CTRL = controls (water-exposed animals); CAS = conspecific alarm substance. Final sample sizes: CTRL + VEH: n = 14 animals; CTRL + CLZ: n = 15 animals; CAS + VEH: n = 14 animals; CAS + CLZ: n = 15 animals.

After CAS exposure, significant main effects of treatment (*p* = 0.0041) and dose (*p* = 0.0023), as well as an interaction effect (*p* = 0.0021), were found for time on top (Figure 4A); CAS decreased time on top (adjusted *p* = 1.07·10^-5^), and CLZ partially blocked this effect. Main effects of treatment (*p* = 2·10^-4^) and dose (*p* = 2.1·10^-4^), as well as an interaction effect (*p* = 0.0044), were found for time on bottom (Figure 4B), and post-hoc tests suggested that CAS increased time on bottom (p = 0.0009) while CLZ blocked this effect. No main effects (*p* > 0.2), nor an interaction effect (*p* = 0.4) were found for absolute turn angle (Figure 4C). A main effect of treatment (*p* = 2.1·10^-4^) and drug (*p* = 2.1·10^-4^), as well as an interaction effect (*p* = 2.3·10^-4^), were found for freezing (Figure 4D); CAS increased freezing (adjusted *p* = 0.0009), and CLZ blocked this effect. No main effects were found for swimming speed (*p* > 0.08), nor were interaction effects found (*p* > 0.3) (Figure 4E).

**Figure 4.**
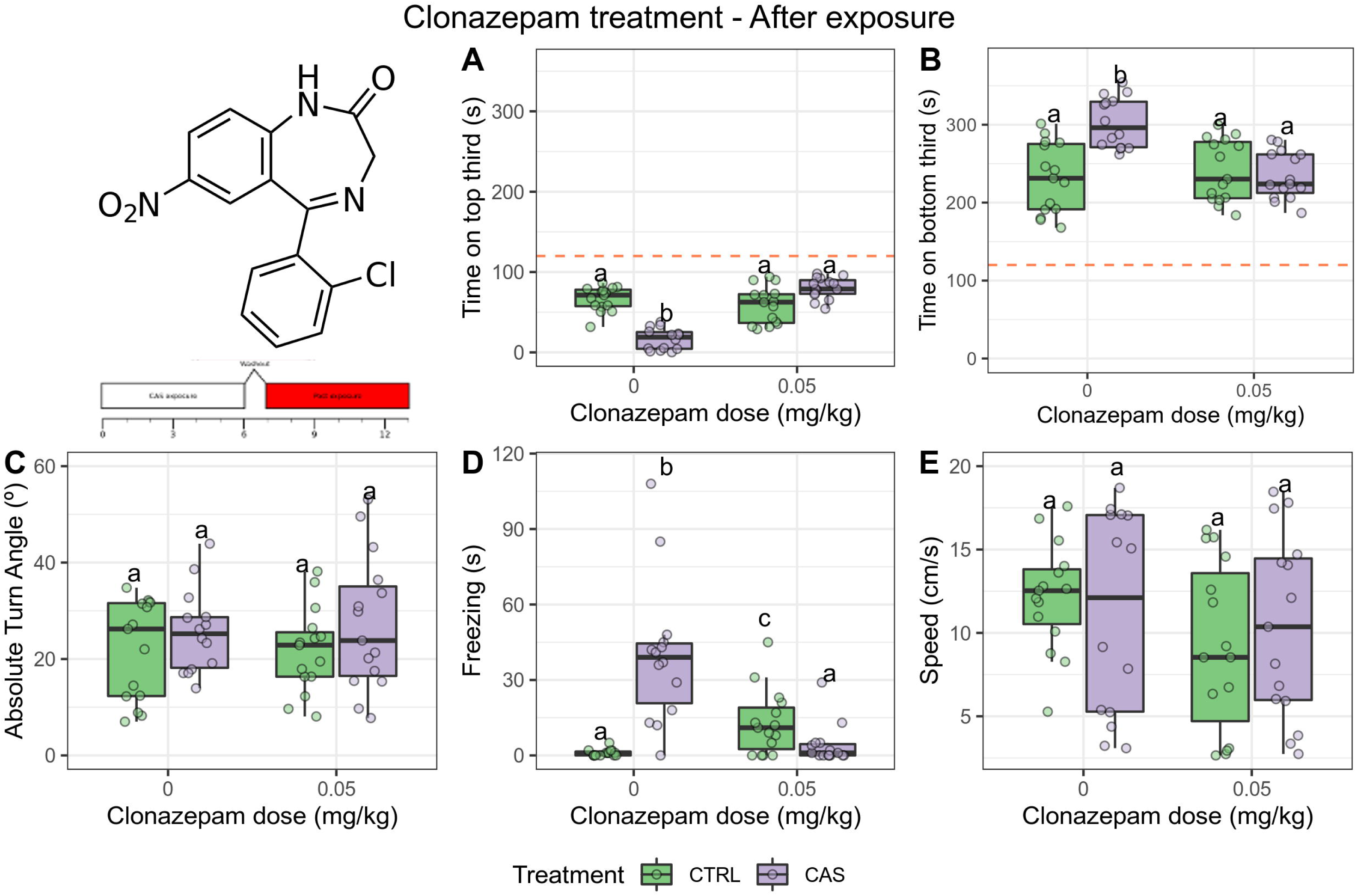
**Clonazepam (0.05 mg/kg) blocks only the CAS-elicited increases in bottom-dwelling after exposure.** (A) Time spent on the top third of the tank; (B) Time spent on the bottom third of the tank; (C) Absolute turn angle; (D) Freezing; (E) Swimming speed. Different letters represent statistical differences at the *p* < 0.05 level; similar letters indicate lack of statistically significant differences. Data are presented as individual data points (dots) superimposed over the median ± interquartile ranges. CTRL = controls (water-exposed animals); CAS = conspecific alarm substance. Final sample sizes: CTRL + VEH: n = 14 animals; CTRL + CLZ: n = 15 animals; CAS + VEH: n = 14 animals; CAS + CLZ: n = 15 animals.

### 3.2. Experiment 2

One outlier was removed from the group exposed to CAS and treated with 2.5 mg/kg fluoxetine. During CAS exposure, significant effects of treatment (*p* = 0.0458), dose (*p* < 0.0001), and interaction (*p* < 0.0001) were found for time on top. Fluoxetine alone increased time on top at 2.5 mg/kg (adjusted *p* = 3.218·10^-5^ vs. control), CAS decreased it (adjusted *p* =0.01132), and fluoxetine blocked the effect of CAS at both doses (Figure 5A). Main effects of treatment (*p* = 0.004) and dose (*p* < 0.0001), but no interaction (*p* =0.6152), were found for time on bottom (Figure 5B); CAS increased time on bottom (adjusted *p* < 0.0236), fluoxetine decreased it at both doses (adjusted *p* < 0.01), and fluoxetine blocked the effect of CAS at the highest dose. A main effect of treatment (*p* = 0.0436), but not dose (*p* = 0.1102) nor interaction (*p* = 0.1148), was found for absolute turn angle (Figure 5C); post-hoc tests suggested that CAS increased absolute turn angle (adjusted *p* = 0.0061 vs. control), and fluoxetine partially (2.5 mg/kg) or fully (25 mg/kg) blocked this effect. A main effect of treatment (*p* < 0.0001) and dose (*p* = 0.0004), as well as an interaction effect (*p* = 0.00002), were found for freezing (Figure 5D); CAS increased freezing (adjusted *p* = 0.0058), and both doses partially blocked this effect. Finally, no effect was found on swimming speed (*p* > 0.39; Figure 5E).

**Figure 5.**
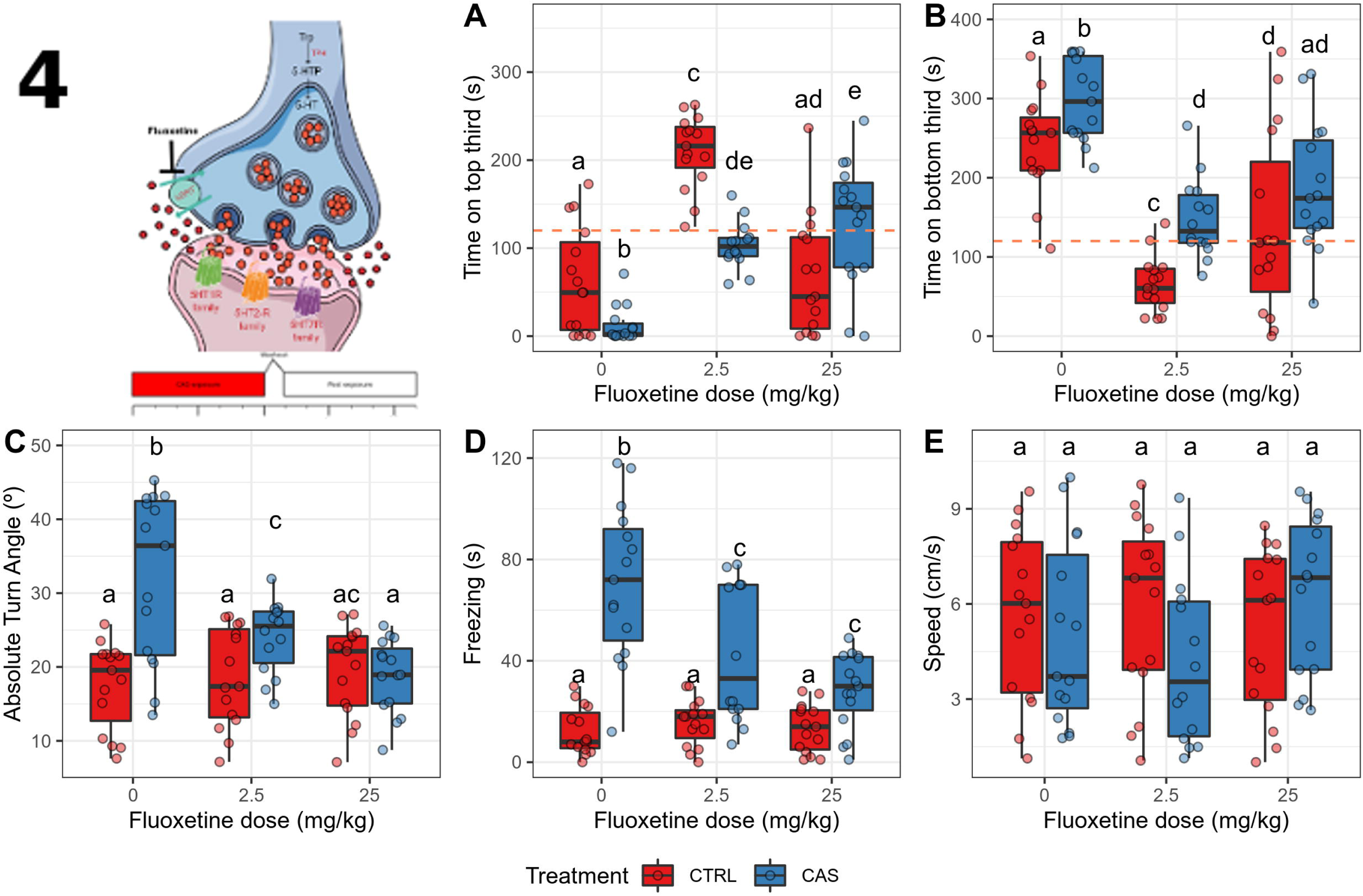
**Acute fluoxetine dose-dependently blocks all the CAS-elicited increases in defensive behavior during exposure.** (A) Time spent on the top third of the tank; (B) Time spent on the bottom third of the tank; (C) Absolute turn angle; (D) Freezing; (E) Swimming speed. Different letters represent statistical differences at the *p* < 0.05 level; similar letters indicate lack of statistically significant differences. Data are presented as individual data points (dots) superimposed over the median ± interquartile ranges. CTRL = controls (water-exposed animals); CAS = conspecific alarm substance. Final sample sizes: CTRL + 0 mg/kg: n = 15 animals; CTRL + 2.5 mg/kg: n = 15 animals; CTRL + 25 mg/kg: n = 15 animals; CAS + 0 mg/kg: n = 15 animals; CAS + 2.5 mg/kg: n = 14 animals; CAS + 25 mg/kg: n = 15 animals.

Significant effects of treatment (*p* = 0.0072) and interaction (*p* = 0.0196) were found for time on top after exposure (Figure 6A). Post-hoc pairwise permutation tests found a difference between control and CAS-exposed animals (adjusted *p* = 1.194 · 10^-5^), an effect that was not blocked by fluoxetine. Similarly, main effects of treatment (*p* = 0.0002), but not a drug (*p* = 0.2914) nor an interaction effect (*p* = 0.5878) were found for time on bottom (Figure 6B), with a significant increase in CAS-exposed animals (all adjusted *p* < 0.001). No effects were found for erratic swimming (*p* > 0.7; Figure 6C). Significant treatment (*p* < 0.0001), dose (*p* = 0.025), and interaction effects (*p* = 0.0014), were found for freezing (Figure 6D), with CAS increasing freezing at all drug treatments, and the highest fluoxetine dose potentiating this effect. No effects were found for swimming speed (*p* > 0.24; Figure 6E).

**Figure 6.**
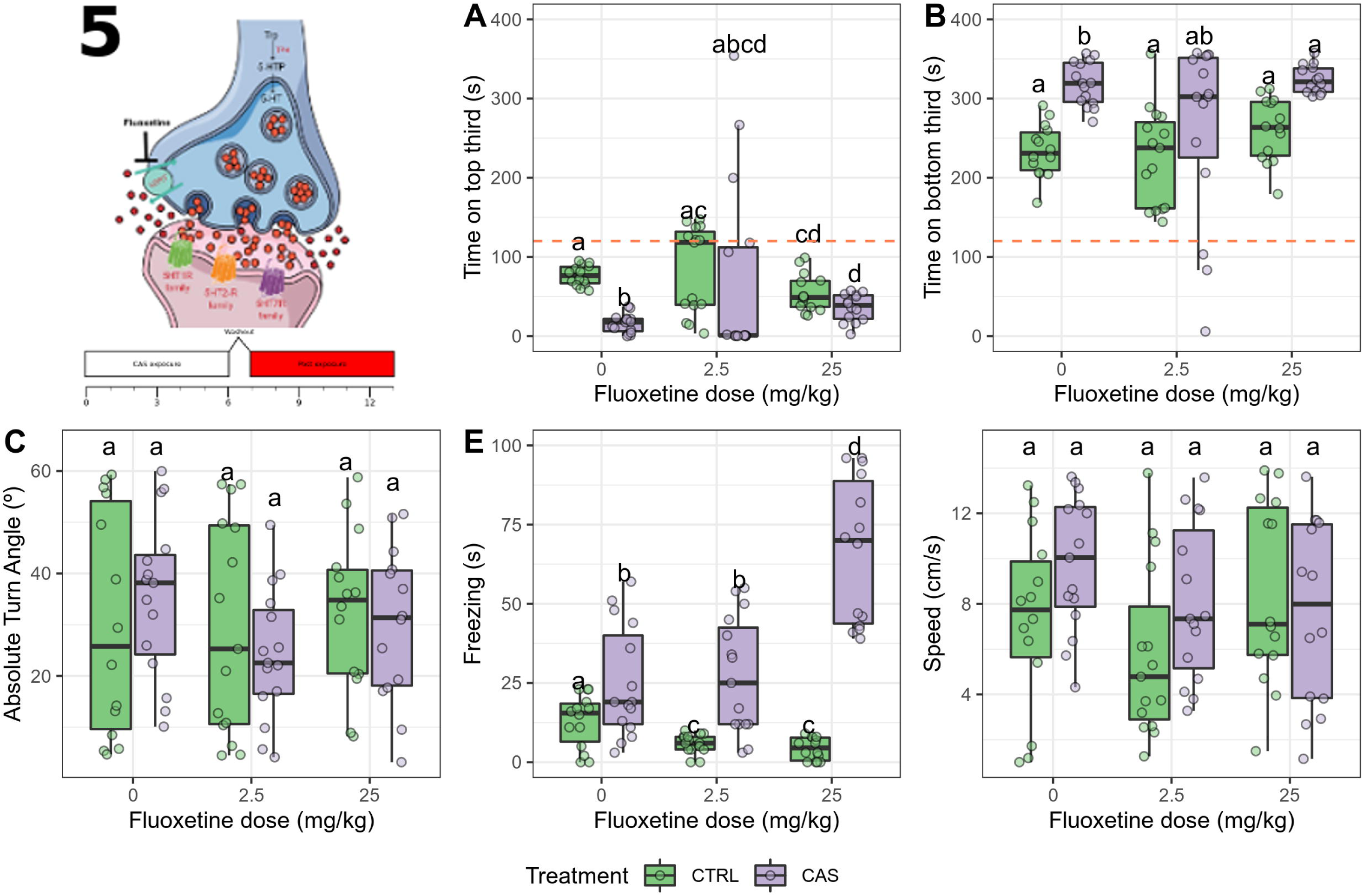
**Acute fluoxetine dose-dependently blocks only the CAS-elicited increases in bottom-dwelling after exposure.**(A) Time spent on the top third of the tank; (B) Time spent on the bottom third of the tank; (C) Absolute turn angle; (D) Freezing; (E) Swimming speed. Different letters represent statistical differences at the *p* < 0.05 level; similar letters indicate lack of statistically significant differences. Data are presented as individual data points (dots) superimposed over the median ± interquartile ranges. CTRL = controls (water-exposed animals); CAS = conspecific alarm substance. CTRL = controls (water-exposed animals); CAS = conspecific alarm substance. Final sample sizes: CTRL + 0 mg/kg: n = 15 animals; CTRL + 2.5 mg/kg: n = 15 animals; CTRL + 25 mg/kg: n = 15 animals; CAS + 0 mg/kg: n = 15 animals; CAS + 2.5 mg/kg: n = 14 animals; CAS + 25 mg/kg: n = 15 animals.

### 3.3 Experiment 3

One outlier was removed from the group exposed to water and treated with vehicle. During CAS exposure, significant effects of treatment (*p* < 0.0001), but not metergoline (*p* = 0.6682) or interaction (*p* = 0.5162), were found for time on top (Figure 7A); post-hoc comparisons suggested that CAS decreased time on top (adjusted *p* = 1.977 · 10^-6^), but metergoline did not block effect. Significant effects of treatment (*p* = 0.0432), but not metergoline (*p =* 0.9518) nor interaction (*p* = 0.4174) were found for time on bottom (Figure 7B), with CAS increasing time on bottom (adjusted *p* = 0.01472) and no effect of metergoline. Significant effects of treatment (*p* = 0.0002), but not metergoline (*p* = 0.6496) nor interaction (*p* = 0.1814), were found for absolute turn angle (Figure 7C), with CAS increasing absolute turn angle (adjusted *p* = 0.02627) and metergoline having no effect. Significant effects of treatment (*p* < 0.0001), but not metergoline (*p* = 0.462) nor interaction (*p* = 0.1922), were found for freezing (Figure 7D); post-hoc tests found significant differences between CAS-exposed animals and controls treated with vehicle (adjusted *p* = 0.02073), but metergoline did not block the effects of CAS on freezing. No effects of treatment (*p* =, metergoline, and interaction (all *p* > 0.1) were found for swimming speed (Figure 7E).

**Figure 7.**
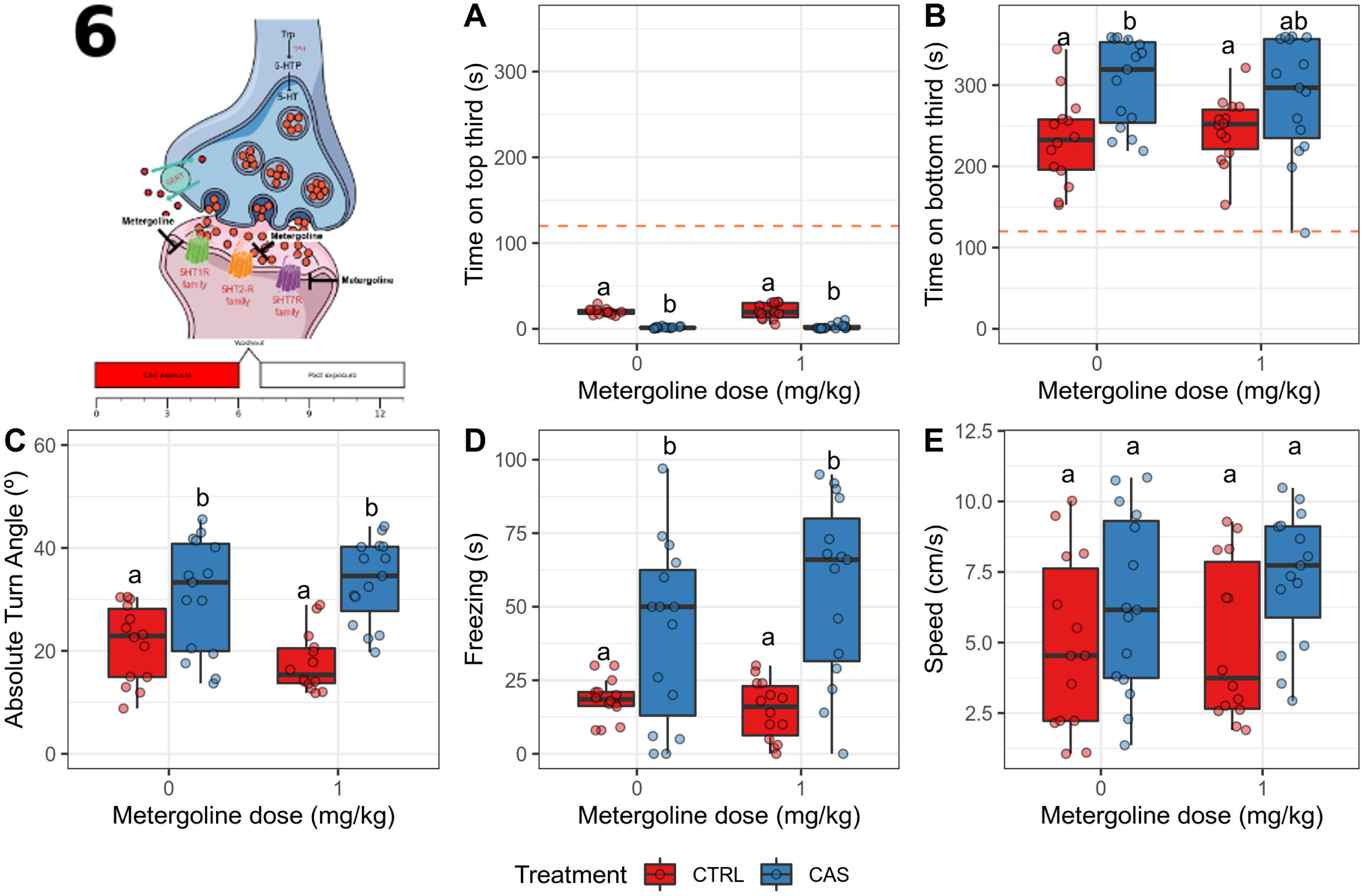
**Metergoline does not block the CAS-elicited increases in defensive behavior after exposure.** (A) Time spent on the top third of the tank; (B) Time spent on the bottom third of the tank; (C) Absolute turn angle; (D) Freezing; (E) Swimming speed. Different letters represent statistical differences at the *p* < 0.05 level; similar letters indicate lack of statistically significant differences. Data are presented as individual data points (dots) superimposed over the median ± interquartile ranges. CTRL = controls (water-exposed animals); CAS = conspecific alarm substance. CTRL = controls (water-exposed animals); CAS = conspecific alarm substance. Final sample sizes: CTRL + VEH: n = 14 animals; CTRL + MET: n = 15 animals; CAS + VEH: n = 15 animals; CAS + MET: n = 15 animals.

After CAS exposure, no main nor interaction effects were found for time on top (all *p* > 0.13; Figure 8A). A dose (*p* = 0.0156) and an interaction (*p* = 0.047) effects were found for time on bottom, but a treatment effect was not found (*p* = 0.827); CAS increased time on bottom (adjusted *p* = 0.0292), an effect that was decreased by metergoline (Figure 8B). No main nor interaction effects were found for absolute turn angle (*p* > 0.33; Figure 8C). A main effect of treatment (*p* = 0.0036), but not drug (*p* = 0.4182), nor interaction (*p* = 0.1508), was found for freezing (Figure 8D); CAS increased freezing (adjusted *p* = 0.0075), and metergoline partially blocked this effect. No main effects were found for swimming speed (*p >* 0.27), but an interaction effect was found (*p* = 0.026); however, post-hoc tests failed to find significant differences between groups (Figure 8E).

**Figure 8.**
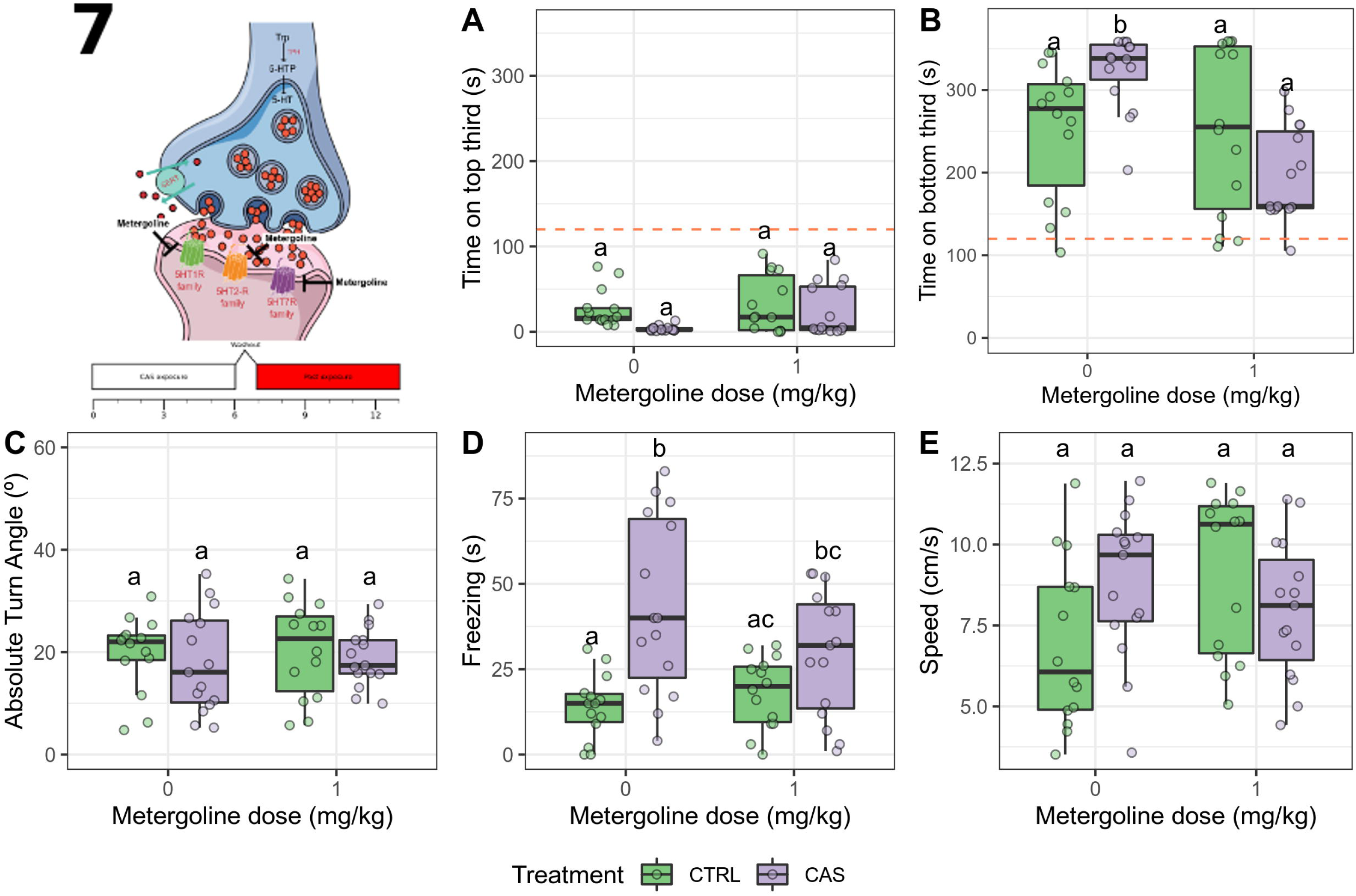
**Metergoline blocks the CAS-elicited increases in bottom-dwellig and freezing after exposure.** (A) Time spent on the top third of the tank; (B) Time spent on the bottom third of the tank; (C) Absolute turn angle; (D) Freezing; (E) Swimming speed. Different letters represent statistical differences at the *p* < 0.05 level; similar letters indicate lack of statistically significant differences. Data are presented as individual data points (dots) superimposed over the median ± interquartile ranges. CTRL = controls (water-exposed animals); CAS = conspecific alarm substance. Final sample sizes: CTRL + VEH: n = 14 animals; CTRL + MET: n = 15 animals; CAS + VEH: n = 15 animals; CAS + MET: n = 15 animals.

### 3.4. Experiment 4

Three outliers were removed from the group exposed to water and treated with vehicle, two from the group exposed to water and treated with pCPA, one from the group exposed to CAS and treated with vehicle, and two from the group exposed to CAS and treated with pCPA. During CAS exposure, a main effect of treatment (*p* < 0.0001), but no drug (*p* = 0.801) nor interaction effects (*p* = 0.5386) were found for time on top (Figure 9A); CAS decreased time on top on both vehicle- and pCPA-injected animals (both ajusted *p* > 0.0025). Likewise, a main effect of treatment (*p* = 0.0022), but no drug (*p* = 0.6496) nor an interaction effects (*p* = 0.1166), were found for time on bottom (Figure 9B); CAS increased time on bottom on both vehicle- and pCPA-injected animals (both adjusted *p* = 0.05). A main effect of treatment (*p* = 0.0002), but not an effect of drug (*p* = 0.8162) nor interaction (*p* = 0.5612), was found for absolute turn angle (Figure 9C); CAS increased absolute turn angle on both vehicle- and pCPA-injected animals (both adjusted *p* < 0.002). A main effect of treatment (*p* < 0.0001), but not an effect of drug (*p* = 0.6822) nor interaction (*p* = 0.4156), was found for freezing (Figure 9D); CAS increased freezing on both vehicle- and pCPA-injected animals (both adjusted *p* < 0.018). No main or interaction effects were found for speed (all *p* > 0.11; Figure 9E).

**Figure 9.**
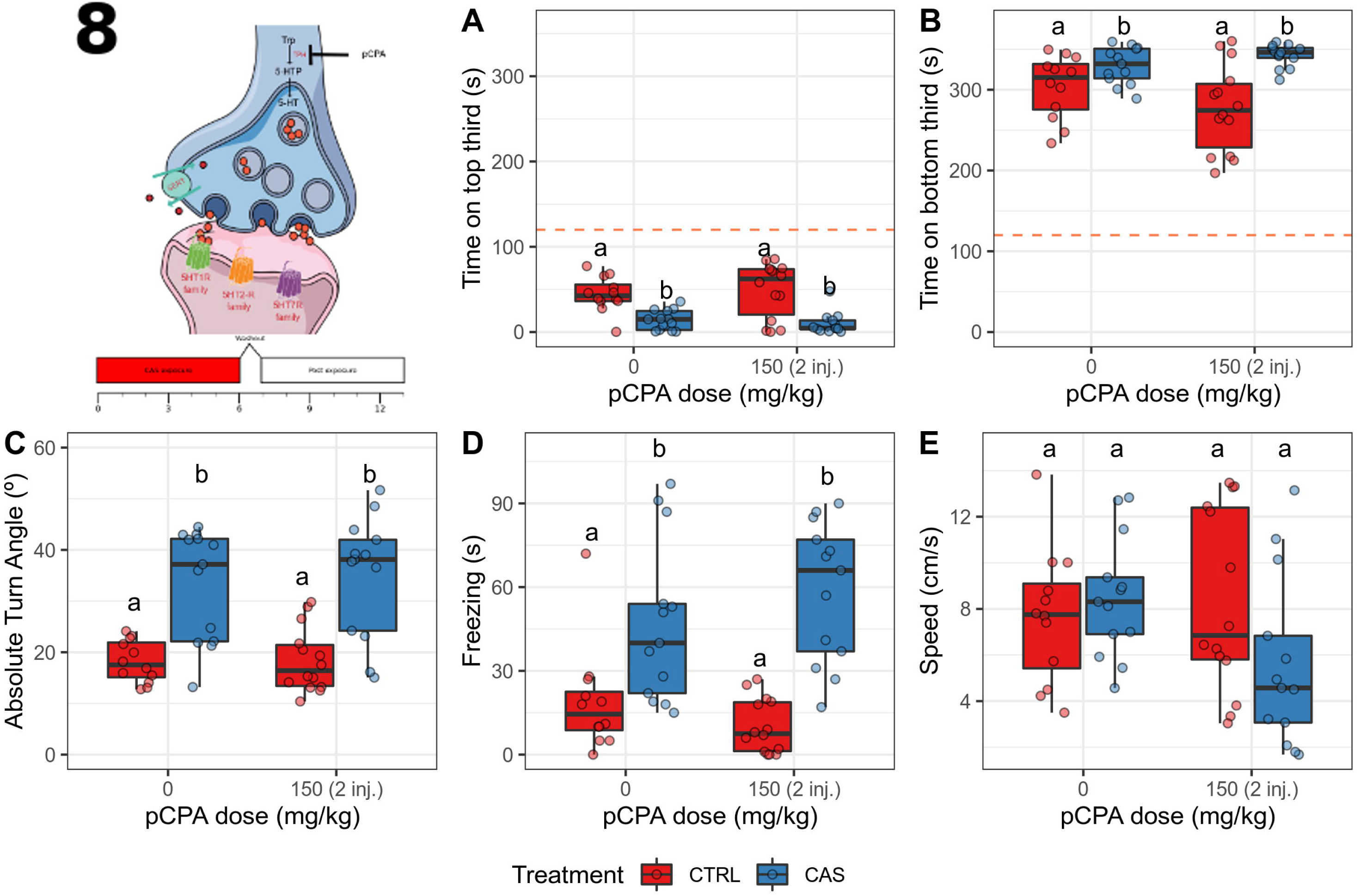
**pCPA does not block the CAS-elicited increases in defensive behavior during exposure.** (A) Time spent on the top third of the tank; (B) Time spent on the bottom third of the tank; (C) Absolute turn angle; (D) Freezing; (E) Swimming speed. Different letters represent statistical differences at the *p* < 0.05 level; similar letters indicate lack of statistically significant differences. Data are presented as individual data points (dots) superimposed over the median ± interquartile ranges. CTRL = controls (water-exposed animals); CAS = conspecific alarm substance. Final sample sizes: CTRL + VEH: n = 12 animals; CTRL + pCPA: n = 13 animals; CAS + VEH: n = 14 animals; CAS + pCPA: n = 13 animals.

After CAS exposure, no main (all *p* > 0.08) or interaction (*p* = 0.1472) effects were found for time on top (Figure 10A). A main effect of drug (*p* < 0.0001), but not of treatment (*p* = 0.4514), nor an interaction effect (*p* = 0.5436), was found for time on bottom (Figure 10B); pCPA decreased time on bottom at both controls and CAS-exposed animals (all adjusted *p <* 0.001). No main (all *p* > 0.3) nor interaction (*p* = 0.6958) effects were found for absolute turn angle (Figure 10C). A main effect of treatment (*p* = 0.0108), but not drug (*p =* 0.1892) nor interaction (*p* = 0.2576), was found for freezing (Figure 10D); CAS increased freezing (adjusted *p* = 0.02278), an effect that was blocked by pCPA (ajusted *p* = 0.01453). No main (all *p* > 0.19) nor interaction (*p* = 0.0954) effects were found for swimming speed (Figure 10E).

**Figure 10.**
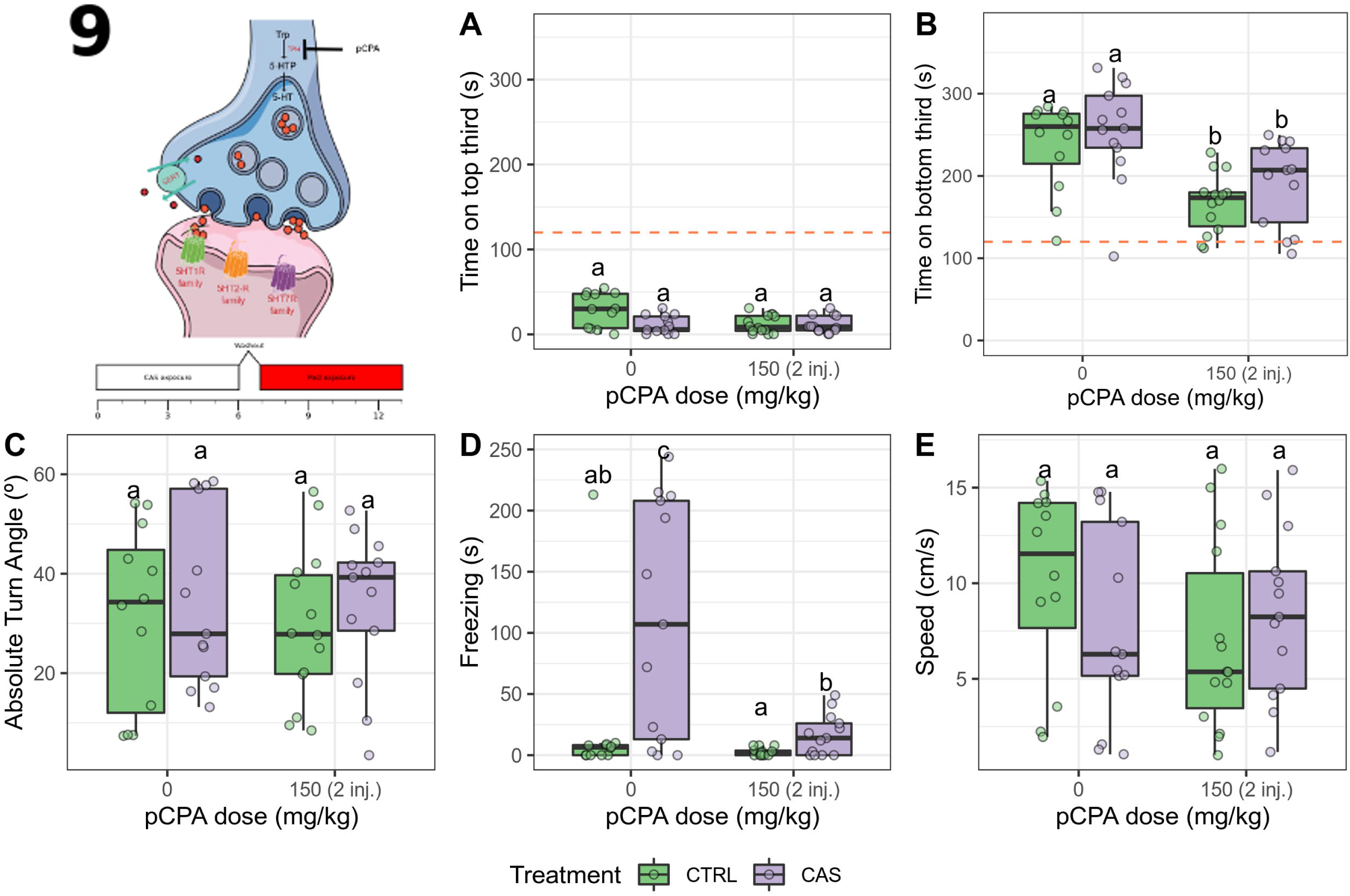
**pCPA blocks the CAS-elicited increases in bottom-dwellig and freezing after exposure.** (A) Time spent on the top third of the tank; (B) Time spent on the bottom third of the tank; (C) Absolute turn angle; (D) Freezing; (E) Swimming speed. Different letters represent statistical differences at the *p* < 0.05 level; similar letters indicate lack of statistically significant differences. Data are presented as individual data points (dots) superimposed over the median ± interquartile ranges. VEH = Vehicle (10% DMSO); pCPA = *para*-chlorophenylalanine; CTRL = controls (water-exposed animals); CAS = conspecific alarm substance. Final sample sizes: CTRL + VEH: n = 12 animals; CTRL + pCPA: n = 13 animals; CAS + VEH: n = 14 animals; CAS + pCPA: n = 13 animals.

### 3.5. Experiment 5

No outliers were removed. After CAS exposure, zMAO activity was reduced in the brain (*Z* = 3.205, *p* = 0.00135; Figure 11).

**Figure 11.**
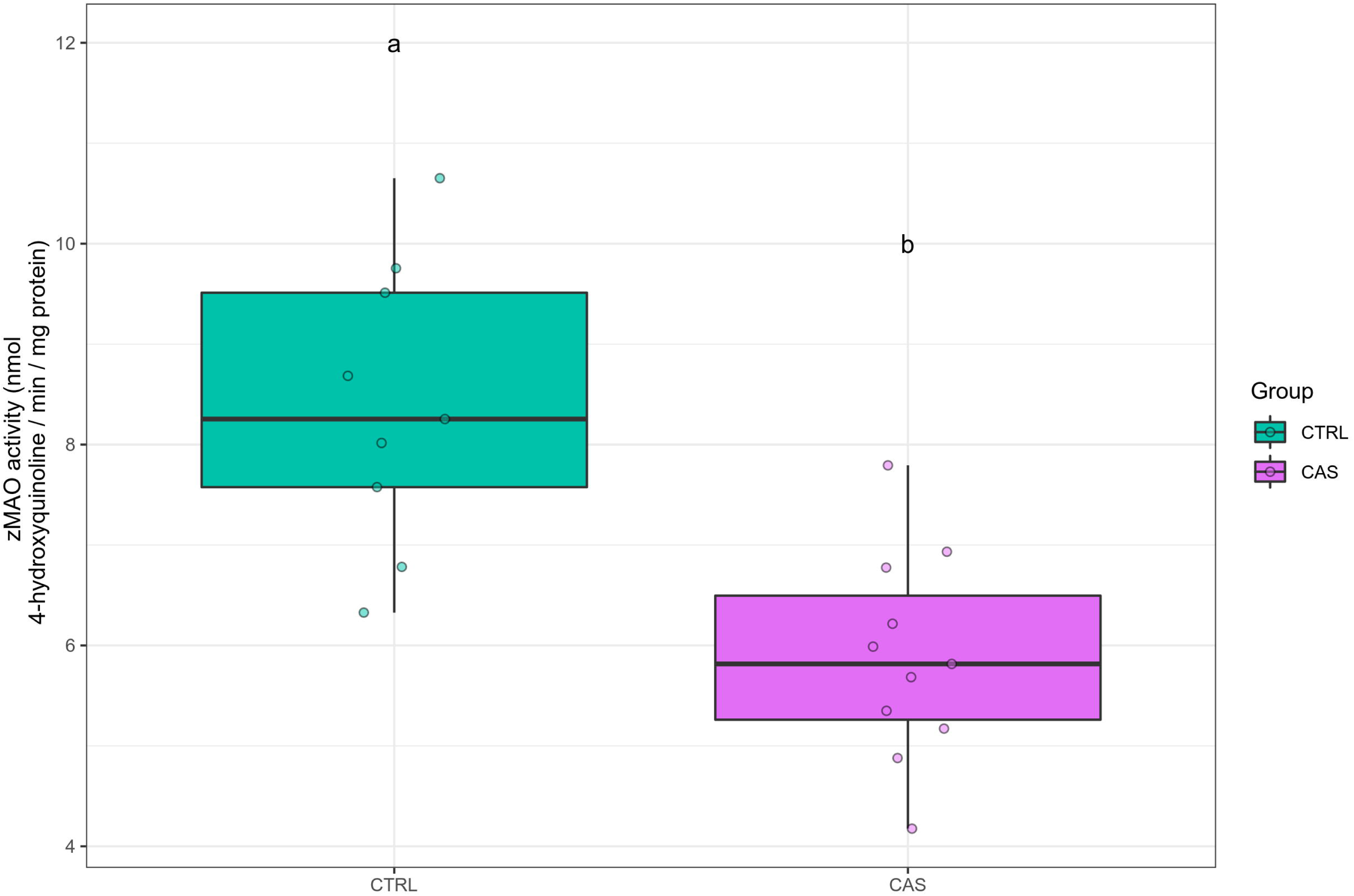
**Conspecific alarm substance (CAS) reduces the activity of monoamine oxidase in the brain after exposure.** Different letters represent statistical differences at the *p* < 0.05 level. Data are presented as individual data points (dots) superimposed over the median ± interquartile ranges. CTRL = controls (water-exposed animals); CAS = conspecific alarm substance. Final sample sizes: CTRL: n = 10 animals; CAS: n = 10 animals.

**Figure 12.**
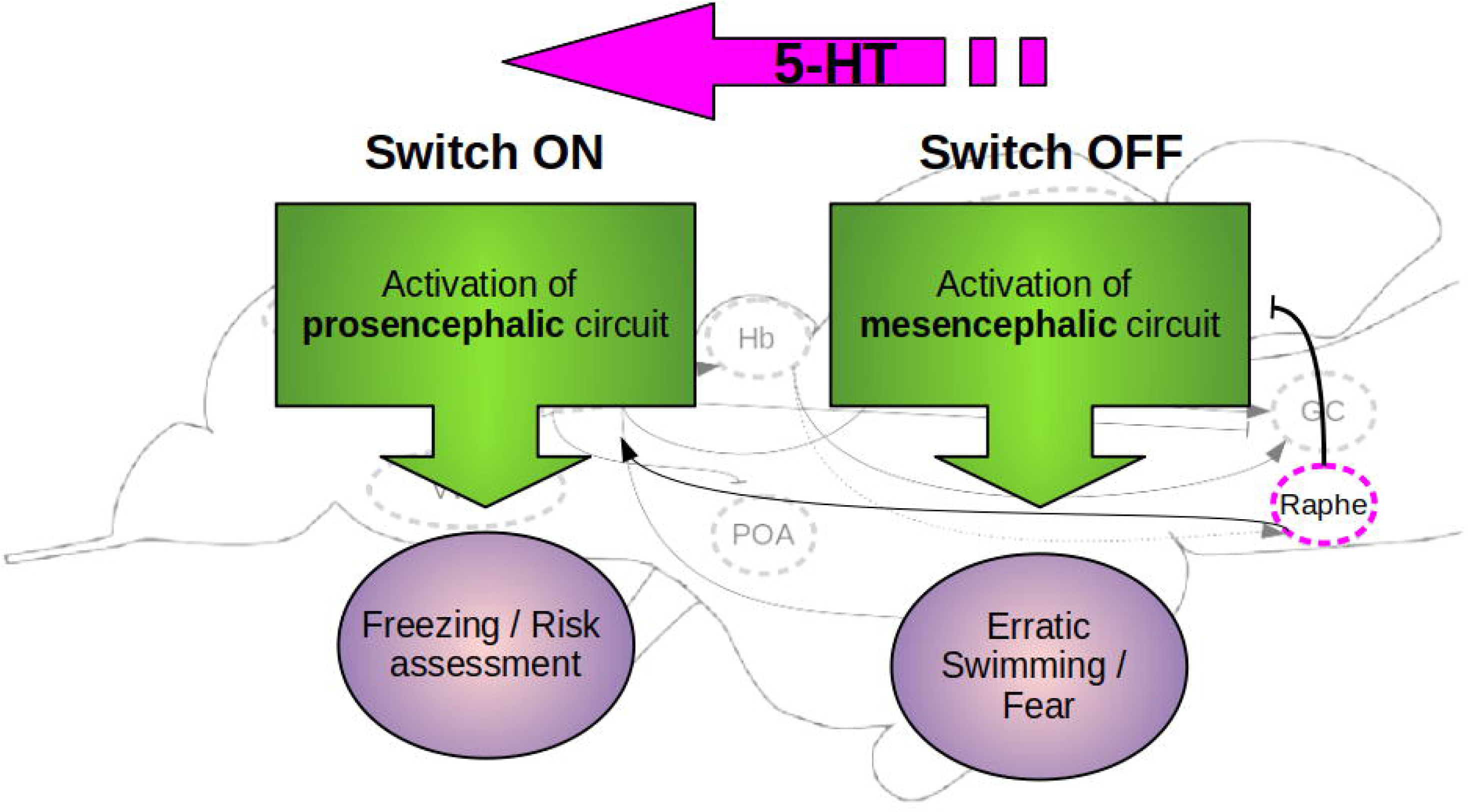
**Hypothetical mechanism of the serotonergic signaling in zebrafish defensive behavior during and after exposure to conspecific alarm substance**. CAS elicits responses dominated by erratic swimming, which decreases as the substance’s concentrations decline. After CAS exposure, the behavioral response is dominated by freezing. Serotonin shifts responding from the first to the second (represented by the purple arrow, as well as by the arrows connecting the raphe to the “switch” green boxes), putatively by switching control from the mesencephalic aversive circuit (“switch OFF”) to the prosencephalic aversive circuit (“switch ON”).

## 4. Discussion

The present work attempted to clarify the role of phasic and tonic serotonin in the alarm reaction of zebrafish during (fear-like behavior) and after (recovery) exposure. We found that clonazepam decreased fear-like behavior, as well as post-exposure behavior, suggesting a good predictive validity of the assay. Moreover, acute fluoxetine decreased fear-like behavior at the highest dose, but increased freezing post-exposure. Metergoline had no effect on fear-like behavior, but blocked the effects of conspecific alarm substance (CAS) on post-exposure behavior; similar effects were observed with pCPA. Finally, CAS was shown to decrease the activity of monoamine oxidase in the zebrafish brain after exposure.

### 4.1. Behavior during and after CAS exposure

In zebrafish, reported behavioral effects of CAS vary widely as a function of timing, extraction method, and whether animals are tested alone or in groups (Maximino *et al.* 2019). When animals are exposed and/or tested alone, as in the present experiments, bottom-dwelling, freezing, and erratic swimming consistently increases *during* exposure (Eachus *et al.* 2017; Nathan *et al.* 2015; Ogawa *et al.* 2014; Maximino *et al.* 2014), but effects *after* exposure are less clear (Quadros *et al.* 2016; Schirmer *et al.* 2013; Nathan *et al.* 2015; Egan *et al.* 2009). In the present experiments, CAS consistently increased bottom-dwelling and erratic swimming during exposure, while after exposure bottom-dwelling and freezing were increased. The first effects were blocked by treatment with the panicolytic drug clonazepam, which nonetheless had a very mild effect on post-exposure behavior. Thus, two components can be elicited by CAS: the first, dominated by erratic swimming, occurs when the substance is present, and the second, dominated by freezing, occurs when the concentrations of CAS decrease.

Observing the context in which defensive behavior, and not only the response topography, is important to understand the function of a specific response. The context in which CAS elicits alarm reactions is akin to a circa-strike defensive situation (Maximino *et al.* 2019), therefore producing freezing and escape reactions that are fear-like; when CAS signals decrease, however (i.e., *after exposure* in the present experiments), the context is akin to a post-encounter defensive situation, eliciting avoidance and freezing behavior (see Perusini and Fanselow 2015 for a discussion on predatory imminence, defensive reactions, and fear vs. anxiety). In these contexts, increases in freezing, for example, can be interpreted as representing two different functions: to escape detection by predators in the first case, and to allow careful vigilance, in the second case.

The differences in behavior during and after exposure are reminiscent of the different types of freezing elicited during and after electrical stimulation of the periaqueductal gray in rodents (Brandão *et al.* 2008). The increased erratic swimming observed during CAS exposure suggest escape and/or avoidance attempts, while the increased freezing observed after CAS exposure suggest a role in risk assessment. The effects of clonazepam also suggest different neurobehavioral states: this drug usually decreases panic attacks, but has small effects on generalized anxiety in human clinical settings (Caldirola *et al.* 2016; Cloos 2005). These results imply good predictive validity, suggesting that behavior during and after CAS can be used to study fear- vs. anxiety-like effects.

### 4.2. Role of phasic serotonin on CAS effects

Fluoxetine, at the highest dose, blocked the alarm reaction (fear-like behavior during CAS exposure) and post-exposure behavior in zebrafish. The results from the lower dose (2.5 mg/kg) are harder to interpret, as they could represent not (partial) blocking, but an additive effect, at least on bottom-dwelling. These results are similar to what was previously observed in the light/dark test, in which fluoxetine (2.5 mg/kg) blocked post-exposure effects on scototaxis, freezing, and erratic swimming (Maximino *et al.* 2014). While the role of phasic increases in serotonergic signaling on acute fear-like responses in zebrafish has not been previously investigated, a higher dose (10 mg/kg) blocked the alarm reaction (i.e., during exposure) in the piauçu *Leporinus macrocephalus* (Barbosa *et al.* 2012). This phasic role of serotonin is likely highly conserved, as serotonin has been shown to decrease responses to aversive odors in *Caenorhabditis elegans* (Li *et al.* 2012; Harris *et al.* 2009). The effects of fluoxetine strongly suggest that serotonin phasically inhibits fear-like behavior in zebrafish, acting as a switch towards risk assessment.

Phasic serotonin has been proposed to modulate fear-like behavior in mammals before (Zangrossi Jr *et al.* 2001; Paul *et al.* 2014; Guimarães *et al.* 2010); the Deakin/Graeff theory suggests a “panic inhibition system” (Paul *et al.* 2014; Silva *et al.* 2019) that inhibits behavioral and sympathoexcitatory responses to these stimuli, and is mediated by serotonergic signaling. The theory proposes a “dual role” for serotonin, increasing anxiety-like responses and inhibiting fear-like responses. A similar mechanism has been proposed for zebrafish based on data regarding serotonergic drugs in anxiety-like behavior (Herculano and Maximino 2014). We propose that, at least in zebrafish, phasic serotonin does not physiologically inhibits fear responses; instead, the inhibitory role of serotonin in fear functions as a “neurobehavioral switch”: as the threatening stimulus ceases, serotonin is released, inhibiting the fear reactions that are now non-adaptive, and initiating careful exploration and risk assessment responses to ensure that the threat is actually over (Figure 11). This is consistent with expectancy value theories, in which serotonin signals represent the expectation of risk/threatening outcomes (aversive expectation values), from which appropriate behavioral strategies can be selected (Amo *et al.* 2014; Cools *et al.* 2011).

A role for the serotonin transporter has also been proposed for the selection of behavior at different levels of a threat imminence continuum: animals with lower expression of the transporter are more cautious and readily show defensive responses under distal threat, while animals with high expression show more defensive responses under proximal threat (Kroes *et al.* 2019). While the expression levels of the serotonin transporter are more associated with controlling serotonergic tone, this protein has been also shown to mediate the increases in serotonin levels after CAS exposure in zebrafish (Maximino *et al.* 2014), suggesting a participation also in phasic signals. Whether serotonin transporter expression levels are associated with the alarm reaction and/or post-exposure behavior in zebrafish is still unknown.

### 4.3. Is there a tonic inhibition of fear-like responses in zebrafish?

The hypothesis that serotonin functions as a “neurobehavioral switch” signal in zebrafish aversive behavior would be strengthened if decreasing the effects of serotonin on its receptors inhibited post-exposure behavior. Indeed, metergoline, which non-specifically blocks 5-HT_1_, 5-HT_2_, and 5-HT_7_ receptors, had no effect on the alarm reaction, but blocked the post-exposure effects of CAS on bottom-dwelling and homebase use; no effect was observed during exposure, suggesting that fear-like responses are not under tonic inhibition. pCPA had similar effects. Nathan et al. (2015) observed that blocking 5-HT_1A_ and 5-HT_2_ receptors potentiates freezing and bottom-dwelling both in the initial moments of exposure and in a “recovery period”; however, during the recovery period animals were still exposed to CAS, the doses which produced effect in Nathan et al. (2015) were higher than reported in other experiments with zebrafish (Maximino *et al.* 2013), and important controls were lacking, making comparison of results difficult.

Further support for this hypothesis is lent by sophisticated experiments made by Amo et al. (2014) using the serotonergic neurotoxin 5,7-DHT. Injection of this neurotoxin in the telencephalon destroyed most serotonergic fibers projecting to it, and led to an inability to learn an active avoidance contingency (Amo *et al.* 2014), suggesting that serotonergic signaling in the telencephalon represents an aversive expectation value. Amo et al. (2014) demonstrated that this circuitry is under the control of projections from the ventral habenula which are not necessary for classical fear conditioning. This suggests that this habenula-raphe-telencephalon pathways do not simply process a fear response, but instead represents expectations values that can be used to inhibit fear when threat is no longer present.

A caveat of the results from metergoline and pCPA experiments is that these drugs also affect other neurotransmitter systems. Metergoline also acts as a non-selective dopamine receptor antagonist (https://www.ebi.ac.uk/chebi/searchId.do?chebiId=CHEBI:64216); although its affinity for 5-HT_2_ receptors is ∼25 times higher than for the dopamine D_2_ receptor, the affinity for 5-HT_1_ receptors is comparable to D_2_ receptors (Dukhovich *et al.* 2004). While pCPA has been reported to produce a selective effect on serotonin levels in zebrafish (Sallinen *et al.* 2009), not altering levels of catecholamines, at the moment we cannot discard the possibility that the treatment used in the present article were not due to changes in these systems. We cannot discard, then, the participation of catecholamines along with serotonin in the effects of these drugs on post-exposure behavior.

In addition to the effects of the manipulations of the serotonergic system on post-exposure behavior, zMAO activity has been shown to be decreased after CAS exposure, which would increase serotonin levels at this moment. Previously, CAS has been shown to increase 5-HT levels in the extracellular fluid of the zebrafish brain 20 min after CAS stress (Maximino *et al.* 2014), and repeated (7 day) exposure to CAS decreases the mRNA levels of the serotonergic genes *pet1* and *slc6a4a* (serotonin transporter)(Ogawa *et al.* 2014). These results suggest that CAS increases serotonergic activity after the stimulus is no longer present, but it is not known whether CAS does so during exposure.

### 4.4. Which receptors are involved?

The present work did not investigate specific receptors which are involved in the alarm reaction in zebrafish. However, a role for 5-HT_1_-, 5-HT_2_-, and 5-HT_7_-like receptors is suggested by the effects of metergoline. The 5-HT_1A_ receptor antagonist WAY 100,635 has been previously shown to block fear-induced analgesia elicited by CAS, but not the increase in anxiety-like behavior in the light/dark test (Maximino *et al.* 2014). At higher doses, however, WAY 100,635 potentiated the effects of CAS both in the early responses (0-8 min) and in the late phase (8-13 min) (Nathan *et al.* 2015). These contradictory results can be explained by differences in exposure methods, as well as differences in behavioral scoring techniques. Methysergide, which non-selectively blocks 5-HT_2A_, 5-HT_2B_, and 5-HT_2C_ receptors, also potentiate the effects of CAS at both times (Nathan *et al.* 2015). However, 5-HT_1A_, 5-HT_2A_, and 5-HT_2C_ receptors in the dorsolateral periaqueductal gray area have been shown to phasically inhibit escape/fear responses in rats (Soares and Zangrossi Jr 2004). While currently it is unknown whether the griseum centrale, the teleostean homolog of the periaqueductal gray area, is involved in fear responses in zebrafish or not, its anatomical position and hodology suggests so (Maximino *et al.* 2019; do Carmo Silva *et al.* 2018a). Thus, 5-HT_1A_ and 5-HT_2_-like receptors appear to be involved in phasic inhibition of fear-like behavior, but so far evidence for a tonic inhibition is lacking.

## 5. Conclusion

The present experiments evidenced two qualitatively different stages of the alarm reaction in zebrafish, one in the presence of the alarm substance, and another when it is no longer present, both sensitive to clonazepam. Results from biochemistry and pharmacological manipulations suggest that phasic and tonic serotonin acts as a neurobehavioral switch towards cautious exploration/risk assessment/anxiety when the aversive stimulus is no longer present. These results refine previous theories on the role of serotonin in anxiety and fear, suggesting new avenues of research.

## List of abbreviations

4-OH: quinoline 4-hydroxyquinoline
5-HT: Serotonin
5,7-DHT: 5,7-dihydroxytryptamine
ANOVA: Analysis of variance
CAS: Conspecific alarm substance
CONCEA: Conselho Nacional de Controle de Experimentação Animal CTRL Control groups
IACUC: Institutional Animal Care and Use Committee
Ibama: Instituto Brasileiro do Meio Ambiente e dos Recursos Naturais Renováveis
IV: Independent variable
MAD: Median absolute difference
pCPA: *para*-chlorophenylalanine
PI: Principal Investigator
ppm: Parts per million
UEPA: Universidade do Estado do Pará
WAY 100,635: *N*-[2-[4-(2-Methoxyphenyl)-1-piperazinyl]ethyl]-*N*-(2-pyridyl)cyclohexanecarboxamide
zMAO: Zebrafish monoamine oxidase

## Acknowledgments

This work was supported by a Conselho Nacional de Desenvolvimento Científico e Tecnológico/CNPq grant to C.M. (#400726/2016-5). M.P.P. was the recipient of a CNPq undergraduate scholarship. D.B.R. is the recipient of a CNPq research productivity grant (#307595/2015-3). The funders had no influence on the study design, the collection, analysis and interpretation of data, as well as writing and submission of this manuscript.

